# Exploratory rearing is governed by hypothalamic MCH cells according to the locus coeruleus

**DOI:** 10.1101/2023.07.19.549648

**Authors:** Cristina Concetti, Paulius Viskaitis, Nikola Grujic, Sian N. Duss, Mattia Privitera, Johannes Bohacek, Daria Peleg-Raibstein, Denis Burdakov

## Abstract

Exploration is essential for survival because it allows animals to gather information about their environment. Rearing is a classic exploratory behavior, during which an animal transiently stands on its hind legs to sample its environment. It is widely observed in common lab conditions as well as in the wild, yet neural signals and circuits underlying this fundamental component of innate behavior remain unclear. We examined behavioral correlates of activity in hypothalamic MCH-producing neurons (MNs) – a recently characterized but still poorly understood neural type – and found that MN activation co-occurs with exploratory rears in mice. Complementary optogenetic and pharmacological manipulations indicated that MN activity selectively promotes rearing via G-protein coupled MCHR1 receptors. Furthermore, we show *in vivo* that activation of the locus coeruleus noradrenergic neurons rapidly inhibits MNs and suppresses rearing through MCHR1-dependent pathways. Overall, these findings define a subcortical neural module which both tracks and controls exploratory rearing.

## INTRODUCTION

Animals innately and spontaneously perform a variety of behaviors fundamental for survival, like locomotion and eating, whose underlying brain circuits are widely studied. Many such behaviors are controlled by hypothalamic neuronal populations^1^ with brain-wide inputs and outputs. Melanin concentrating hormone-producing neurons (MNs) are a genetically-defined, brain-wide projecting population in the lateral hypothalamus (LH) that release the MCH peptide as a neurotransmitter acting on brain-wide distributed MCH G-protein coupled receptors^2–6^. They are classically known to be involved in energy homeostasis^7–10^ and sleep regulation^11^, but have more recently been found to intervene also in learning and plasticity phenomena^12–18^, and in the stabilization of hippocampal theta rhythm^19^, which is associated to exploration and learning^20, 21^. The activity profile of MNs has been investigated in relation to neutral stimuli^12, 13, 22^, appetitive stimuli^23, 24^, and aversive stimuli^25^, but their involvement in self-paced behaviors is only partly understood^24^.

Unsupported rearing is an exploratory behavior displayed by rodents and other mammals by lifting their front body and supporting themselves only on their hind legs^26–28^. This behavior is thought to allow them to reach information streams that surpass those available at ground level^27^. Another advantage is related to defence when animals gather information about potential threats^29^: rearing behavior enables an animal to fulfil its exploratory motivation from a distance, without having to fully commit or engage in overt actions^27^. In the context of rearing related to spatial sampling of the environment and updating spatial information^30–32^, it has been found to correlate with the activity of hippocampal neurons^33^. Rearing has been shown to be reduced by acute stressors^26^, which are known to increase hippocampal activity^34, 35^ and recent work has shown that optogenetic activation of the locus coeruleus (LC) suppresses rearing^36^, in agreement with past observations that rearing is sensitive to the overall level of arousal^26, 37^. MNs rapidly turn on and off during exploration^12, 13, 22^, are inhibited by noradrenaline^38^, and the LH is a site of LC innervation^39^. However, whether these rapid changes in MN activity coincide with rearing is unknown, and MNs’ causal links to rearing remain unprobed.

Here, we investigated relationships between endogenous MN activity and a set of fundamental behaviors - including rearing - that mice can display at their own pace and by their own initiative, in a neutral environment. By combining neural recordings, pharmacology, and optogenetics with a machine learning-based classification of behaviors, we found that MNs report and control rearing behavior, and serve as a downstream effector of LC-noradrenergic influence on rearing.

## RESULTS

### MNs report rearing behavior

To investigate the natural activity of MNs during various self-paced behaviors, we performed fibre photometry using the calcium indicator GCaMP6s under the *Pmch* promoter (Fig. 1A) while video tracking mouse behavior in an open field (Fig. 1B). We used a machine learning behavioral classifier tool^40^ based on a convolutional neural network to identify self-paced behavior (Fig. 1C). We defined five fundamental self-paced behaviors based on specific criteria (see Methods): rearing, grooming, immobility, locomotion and turning. The output of the classifier was then used to identify neuronal activity simultaneous with self-paced behavioral events. An additional behavior, licking, was recorded in a separate chamber equipped with a capacitive touch sensor to detect licking from a spout through which liquid food was delivered. Fig. 1D-I shows examples of MN activity corresponding to the behaviors. Using this approach, we were able to analyse behavior-associated neuronal activity with high accuracy and temporal resolution (∼90% and 3 Hz).

**Figure 1:**
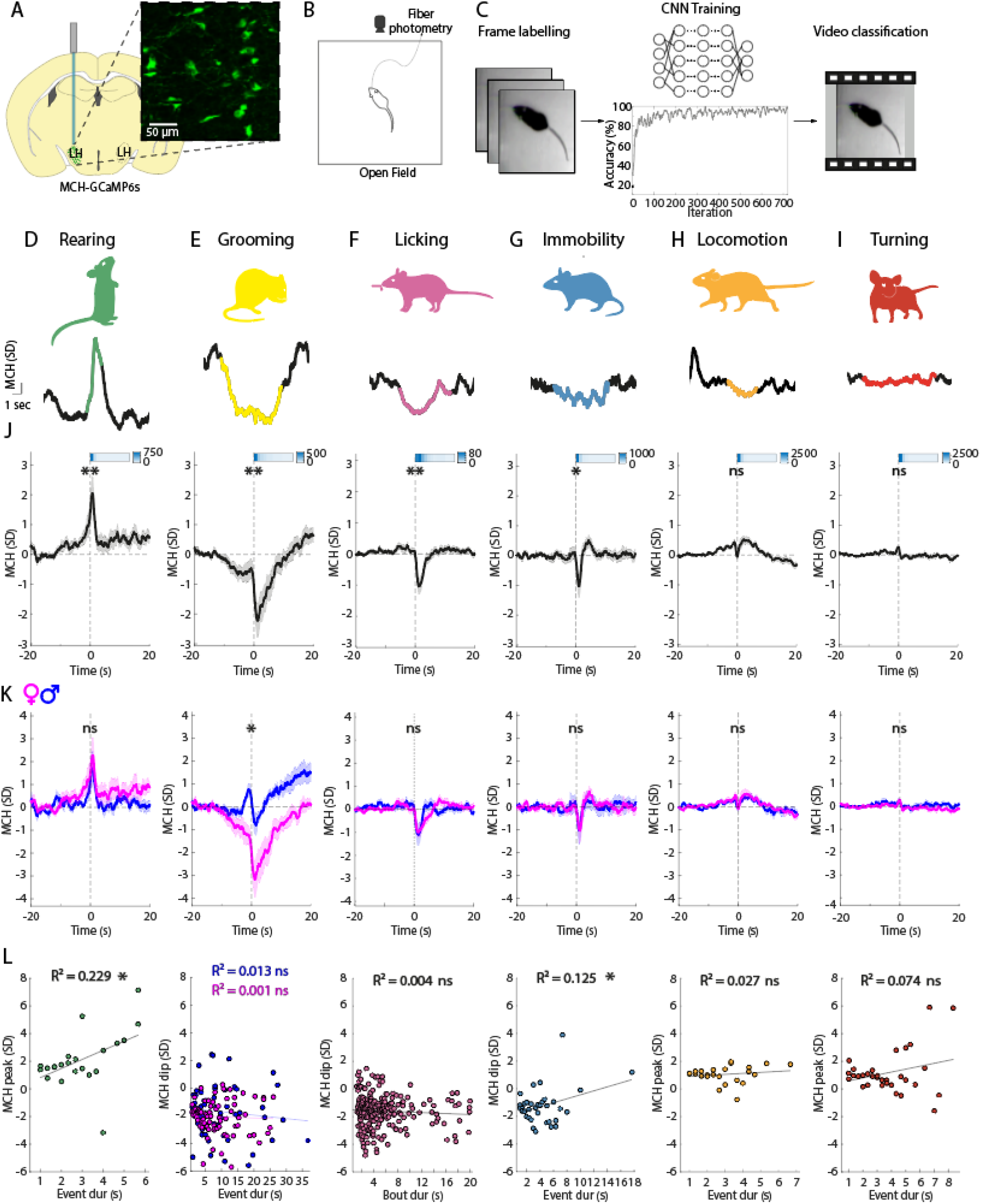
MN activation patterns aligned to initiation of self-paced behaviors. A) Targeting scheme and expression of GCaMP6s in MNs. B) Schematic of the open field experimental paradigm, with video tracking and fibre photometry recording. C) Workflow for behavioral classification using a Convolutional Neural Network (see Results and Methods for details). D-I) Examples of MN activity simultaneous to various self-paced behaviors. J-L) Behavior-associated MN activity as recorded with fibre photometry in the open field. J) Each plot is an average across recording sites and behavioral events. Rearing ** p = 0.0017, grooming ** p = 0.0013, licking ** p = 0.0011, immobility * p = 0.0367, locomotion ns p = 0.0927, turning ns p = 0.4076; one-sample t-tests; n = 26 recording sites from 16 mice. The bars on the top-right of each graph are heatmaps representing the temporal distribution of behavioral events (heatmap units are raw numbers of events). Data are presented as mean ± SEM. K) Same data as in J, separately for males and females. Rearing ns p = 0.7021; grooming * p = 0.0108; licking ns p = 0.7809; immobility ns p = 0.9440; locomotion ns p = 0.9664; turning ns p = 0.9283; n = 10 recording sites from 6 male mice, n = 16 recording sites from 10 female mice; two sample unpaired t-test. L) correlation between the amplitude of MN activity (positive amplitude for behaviors associated with a positive or no deflection, and negative amplitude for those associated with a negative deflection) and event duration of the corresponding behavior. Rearing * p = 0.024, R^2^ = 0.229; grooming ns p = 0.275 R^2^ = 0.013; licking ns p = 0.306 R^2^ = 0.004; immobility * p = 0.038 R^2^ = 0.125; locomotion ns p = 0.442 R^2^ = 0.027; turning ns p = 0.125 R^2^ = 0.074. The black line represents linear regression. ns, p > 0.05; *, p < 0.05; **, p < 0.01.

MN activity showed different profiles across the behavioral variables (Fig. 1J). MN activity significantly increased during rearing behavior, significantly decreased during grooming, licking and immobility, but was unchanged during locomotion and turning. Next, to further analyse the relationship between MN activity and behaviors, we investigated whether there are sex differences, by comparing the amplitude of MCH signals in male and female mice (Fig. 1K). The data showed a significant sexual dimorphism only in the case of grooming, but not for other behaviors. We also analysed whether there is correlation between the amplitude of MN activity and event duration of the corresponding behavior (Fig. 1L). The data showed a significant correlation between the duration of rears and the amplitude of rearing-associated MCH activity. The association between the immobility-associated MCH signal reduction and immobility event duration was also significant, but weaker. Together, these data show that, during spontaneous behavioral sequences, MN activity is increased during self-paced rears in both male and female mice, and rearing-associated MCH amplitude positively correlates with the duration of rears.

### MNs control rearing behavior

To investigate whether and how the endogenous MCH activity influences the tested behaviors, we administered an antagonist of MCH-R1 (the only MCH receptor in mice^41^), SNAP-94847 (20 mg/kg) or vehicle, via i.p. injection before testing (Fig. 2A). SNAP-94847 did not affect center-border preference (a measure of anxiety-like behavior) nor locomotor activity (Fig. 2B-C). However, mice treated with SNAP-94847 showed a significant decrease in time spent rearing, compared to vehicle-injected mice (Fig. 2D). No other behavior was affected by treatment with the MCH-R1 antagonist (Fig. 2E-I). These data suggest that the MCH system is involved in promoting rearing behavior, and that this effect is not due to potential locomotion or anxiety-related effects of SNAP.

**Figure 2:**
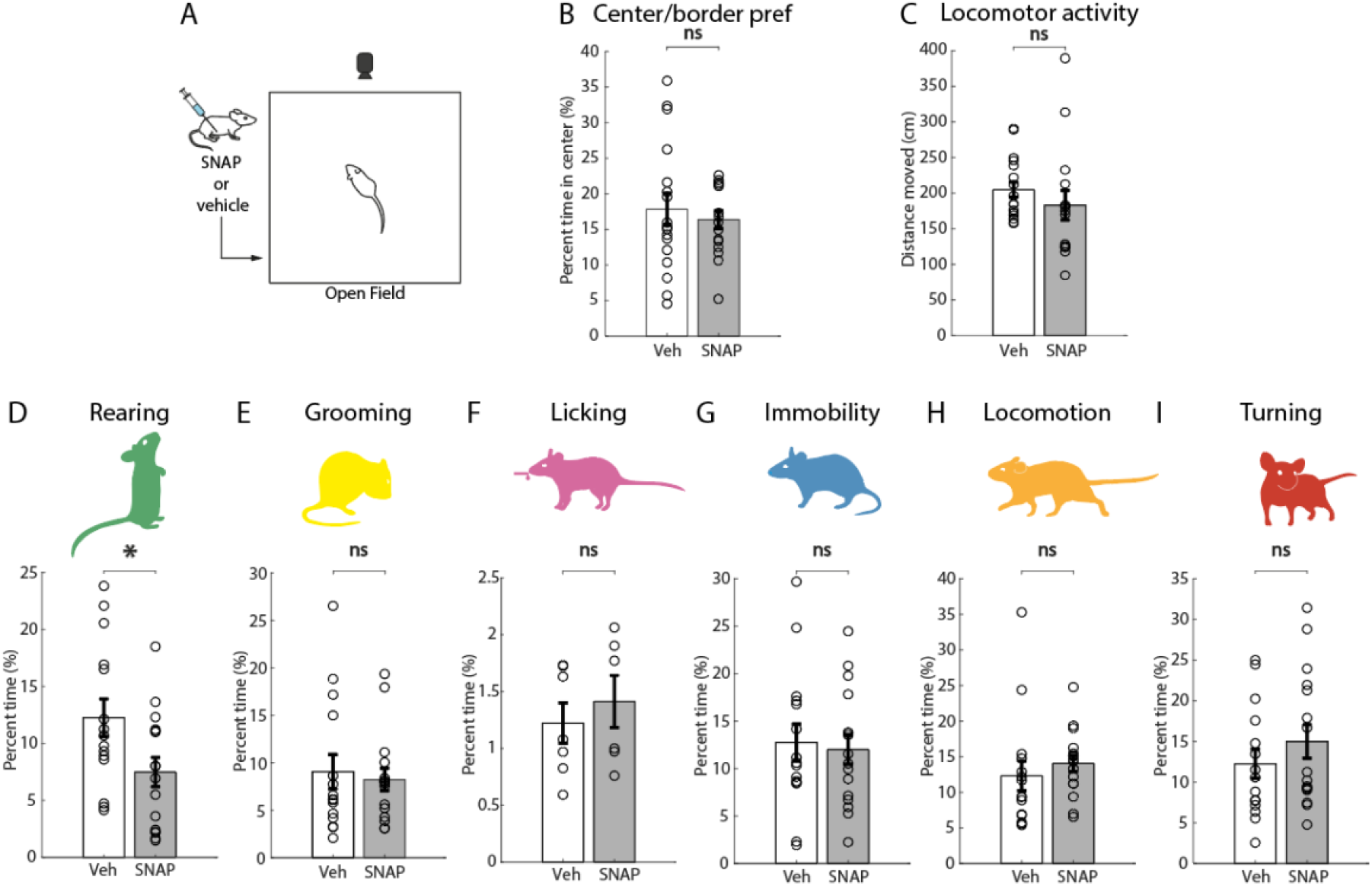
Effects of MCHR1 antagonist SNAP on self-paced behaviors. A) Schematic representation of the experimental paradigm. B-C) Effect of treatment with SNAP (20 mg/kg) vs vehicle on center preference (*ns* p = 0.5903), an indicator of anxiety-like behavior, and locomotor activity in the open field (p = 0.3386; unpaired t-test; n = 15 vehicle mice, 16 SNAP mice). D-I) Effect of treatment with SNAP (20 mg/kg) vs vehicle on time spent performing each behavior. Rearing *\** p = 0.0223; grooming *ns*, p = 0.9249; licking *ns* p = 0.5211; immobility *ns* p = 0.9110; locomotion *ns* p = 0.4626; turning *ns* p = 0.4574; unpaired t-test, n = 15 vehicle mice, 16 SNAP mice for all behaviors except licking where n = 7 vehicle mice and 6 SNAP mice. Data are shown as mean ± SEM. ns, p > 0.05; *, p < 0.05; **, p < 0.01.

In view of these data, we hypothesised that MN activation may selectively increase rearing behavior. To test this, we injected a Cre-dependent excitatory opsin, ChrimsonR, in the LH of MCH-Cre^+^ mice^42^ (Fig. 3A). We then recorded self-paced behaviors while delivering bilateral laser light to the lateral hypothalamus of MCH-ChrimsonR-expressing and control mice (Fig. 3B). Optogenetic activation of MNs did not affect anxiety-like behavior nor locomotor activity (Fig. 3C-D). Rearing levels before optogenetic stimulation were not different between groups (Fig. 3E). Rearing was significantly increased by optogenetic stimulation of MNs compared to controls (Fig. 3D), but no other behavior was affected (Fig. 3E-I).

**Figure 3:**
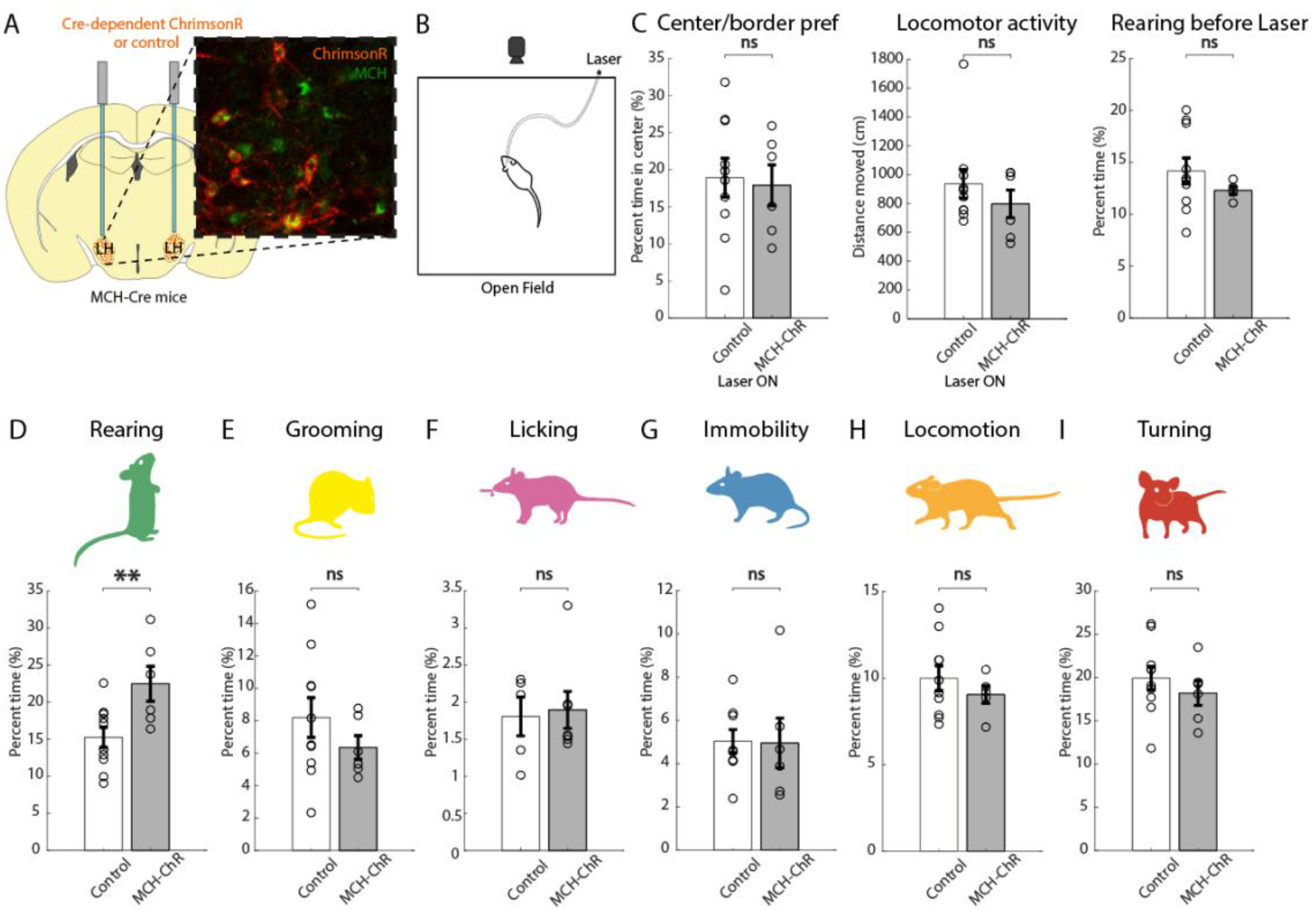
Effect of optostimulation of MNs on self-paced behaviors. A) Targeting scheme and expression of the excitatory opsin ChrimsonR in MNs. B) Schematic representation of the experimental paradigm (stimulation 635 nm, 30 Hz, 10 ms ON, 7 mW, 3 min OFF – 3 min ON – 3 min OFF). C) Effect of laser light stimulation in ChrimsonR-expressing mice versus control mice on center border preference (*ns* p = 0.8019), locomotor activity (*ns* p = 0.3692) during laser stimulation, and rearing levels before laser stimulation (*ns* p = 0.8116; unpaired t-test). D-I) Effect of laser light stimulation in ChrimsonR-expressing mice versus control mice on time spent performing each behavior. Rearing *\*\** p = 0.0062; grooming *ns* p = 0.8521; licking *ns* p = 0.2286; turning *ns* p = 0.7925; locomotion *ns* p = 0.7965, immobility *ns* p = 0.5262; unpaired t-test; n = 5 ChrimsonR-expressing mice and 10 control mice, except for licking where n = 7 ChrimsonR-expressing mice and 5 control mice. Data are shown as mean ± SEM. ns, p > 0.05; *, p < 0.05; **, p < 0.01.

The time spent performing a given behavior is a result of behavioral event frequency (a measure of behavior initiation), and duration (a measure of behavior maintenance). To gain more understanding into how the MCH system regulates rearing, we therefore examined these finer elements of rearing temporal microstructure (Fig. 4A). We found that both frequency and duration of rears are decreased by SNAP compared to vehicle treatment (Fig. 4B). In turn, optogenetic activation of MNs results in a significant increase in both frequency and duration (Fig. 4C). Finally, given that MNs are known to express several neurotransmitters in addition to MCH^19, 43, 44^, we asked whether the rearing-increasing effects of MCH cell optostimulation require MCH receptors. We found that SNAP prevented MCH cell optostimulation from increasing rearing behavior (Fig. 4D). This suggests that the ability of MNs to control rearing requires MCH receptors. Taken together, these results suggest that the MNs and MCH receptors are specific modulators of rearing behavior, controlling both its initiation and maintenance.

**Figure 4:**
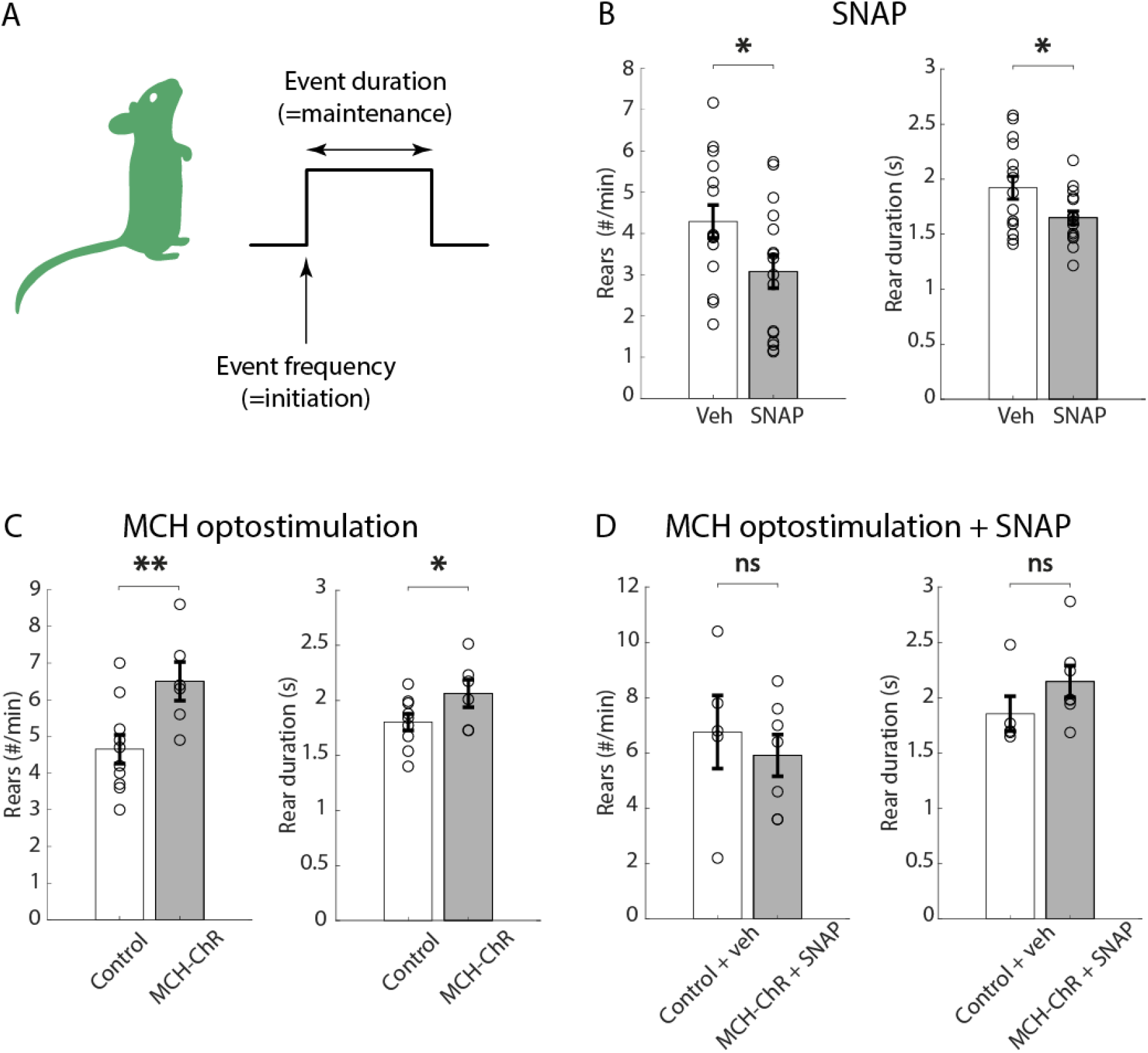
Further dissection of effects of MCH system manipulations on rearing behavior. A) Graphical illustration of behavioral microstructure studied. B) Effect of treatment with SNAP (20 mg/kg) vs vehicle on rear frequency (*\** p = 0.0397) and duration (*\** p = 0.0279; unpaired t-test, n = 15 vehicle mice and 16 SNAP mice). C) Effect of laser light stimulation in ChrimsonR-expressing mice versus control mice on rear frequency (*\*\** p = 0.0063) and duration (*\** p = 0.0417; unpaired t-test; n = 5 ChrimsonR-expressing mice and 10 control mice; stimulation 635 nm, 30 Hz, 10 ms ON, 7 mW, 3 min OFF – 3 min ON – 3 min OFF). D) Effect of simultaneous light stimulation and treatment with SNAP (20 mg/kg) vs vehicle in ChrimsonR-expressing and control mice, on rear frequency (*ns* p = 0.2828) and duration (*ns* p = 0.8957; unpaired t-test; n = 6 ChrimsonR-expressing mice and 5 control mice; stimulation 635 nm, 30 Hz, 10 ms ON, 7 mW, 3 min OFF – 3 min ON – 3 min OFF). Data are shown as mean ± SEM. ns, p > 0.05; *, p < 0.05; **, p < 0.01.

### The MCH system as an effector of noradrenergic influence on rearing

Stressful/threatening environments can suppress rearing^26–28^. Under stressful circumstances, locus coeruleus noradrenergic neurons are thought to mediate central and peripheral responses to stress^45–48^ and LC activation in the open field can suppress rearing^36^. Therefore, we sought to investigate whether LC-noradrenergic neurons inhibit MNs *in vivo,* and hypothesized that optogenetic manipulation of LC-noradrenergic neurons may affect rearing by modulating MNs. To do this, we injected the Cre-dependent optogenetic activator ChrimsonR in the LC of DBH-iCre^+^ mice, a mouse line expressing the Cre recombinase specifically in LC-noradrenergic neurons^49^. In the same mice, we injected the MCH-promoter dependent calcium activity indicator GCaMP6s in the lateral hypothalamus (Fig. 5A). We functionally confirmed the effectiveness of optogenetic stimulation of LC DBH-ChrimsonR neurons by observing pupil dilation in response to the LC optostimulation (Fig. 5B) ^50–52^. Next, we recorded MCH-GCaMP6s cell activity in the open field, while optostimulating the LC noradrenergic neurons (Fig. 5C). At the onset of laser illumination, both MN activity and rearing significantly decreased in ChrimsonR-expressing mice, but not in control mice (Fig. 5D-F).

**Figure 5:**
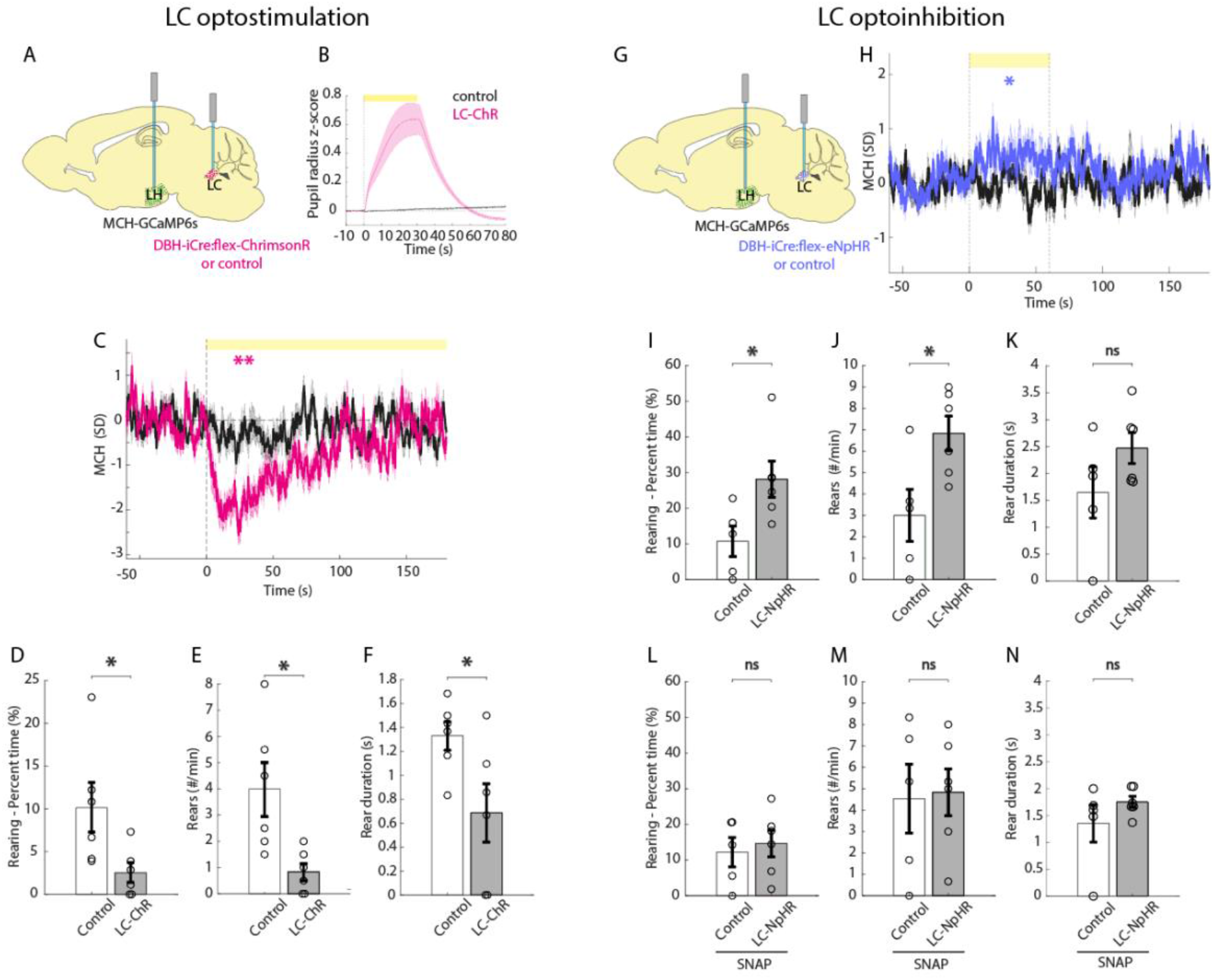
Effects of bidirection optogenetic manipulations of LC-noradrenergic neurons on MNs and rearing. A) Schematic for targeting GCaMP to MNs and ChrimsonR to LC-noradrenaline cells. B) Pupil diameter response to laser illumination of LC in LC-ChrimsonR and control mice. C) Fibre photometry response of MNs to laser illumination of the LC in LC-ChrimsonR mice (*\*\** p = 0.001, magenta) and control mice (*ns* p = 0.6247, black; one-sample t-tests on baseline-subtracted average activity during the first 60 sec of laser illumination; n = 12 recording sites from 6 mice for LC ChrimsonR, n = 12 recording sites from 6 mice for controls; stimulation 635 nm, 30 Hz, 10 ms ON, 7 mW, 3 min OFF – 3 min ON – 3 min OFF). D-F) Rearing behavior during MN activity inhibition caused by optostimulation of LC-noradrenergic neurons in LC-ChrimsonR mice compared to control mice. Rearing percent time *\** p = 0.0182; rear frequency *\** p = 0.0071; rear duration *\** p = 0.0201; unpaired t-test; n = 6 LC-ChrimsonR expressing mice and 6 control mice; stimulation 635 nm, 30 Hz, 10 ms ON, 7 mW, 3 min OFF – 3 min ON – 3 min OFF. G) Targeting schematic of MCH-dependent GCaMP in the LH and Cre-dependent inhibitory opsin eNpHR in the LC of DBH-iCre mice. B) Fibre photometry response to laser illumination of the LC and simultaneous stress cue in LC-eNphR mice (*\** p = 0.0484) and control mice (*ns* p = 0.7465; one-sample t-tests; n = 6 eNpHR-expressing mice and 5 control mice; stimulation 589 nm, CW, 7 mW, 1 min ON followed by 1.5 min OFF x 6 times). I-K) Rearing behavior following laser illumination of the LC and simultaneous stress cue in LC-eNpHR and control mice. Rearing percent time *\** p = 0.0151; rear frequency *\** p = 0.0121; rear duration *ns* p = 0.0796, n = 6 eNpHR-expressing mice and 5 control mice. L-N) Rearing behavior in SNAP treated mice following acute stress and laser illumination of the LC in LC-eNpHR and control mice. Rearing percent time *ns* p = 0.3343; rear frequency *ns* p = 0.4386; rear duration *ns* p = 0.1328; unpaired t-test; n = 6 eNpHR-expressing mice and 5 control mice; stimulation 589 nm, CW, 7 mW, 1 min ON followed by 1.5 min OFF x 6 times. Data are shown as mean ± SEM. ns, p > 0.05; *, p < 0.05; **, p < 0.01.

Next, we injected the Cre-dependent silencing opsin eNpHR into the LC and MCH-promoter-driven GCaMP6s in the LH of DBH-iCre^+^ mice (Fig. 5G). To avoid a floor effect (due to low LC activity) while studying the effects of the LC optosilencing, we employed a paradigm involving an acute stressor. We first subjected mice to fear conditioning, where a tone was associated with a foot shock, then played this tone (now serving as stress-inducing cue) simultaneously with optosilencing. The LC optosilencing increased MN activity (Fig. 5H) and increased rearing in eNpHR-expressing, but not in control, mice (Fig. 5I-K). The MCH-R1 antagonist SNAP blocked the effect of the LC optosilencing in rearing (Fig. 5L-N). Interestingly, in addition to rearing, the LC optomodulation altered some other behaviors, like grooming and locomotion but not on immobility and turning (Fig. S1A-H), and these changes were largely unaffected SNAP (Fig. S1I-L). Additionally, the LC optostimulation decreased center preference and locomotor activity (Fig. S2A-B) while LC optosilencing increased center preference and created a trend towards increased locomotor activity (Fig. S2C-D), as expected^36, 45–48, 53–55^, and the latter effect was not changed by MCHR1-antagonist SNAP (Fig. S2E-F).

In summary, we found that optostimulation of LC-noradrenergic neurons inhibits MN activity and rearing. On the other hand, optoinhibition of LC-noradrenergic neurons disinhibits MNs and causes an increase in rearing behavior, which is abolished by treatment with the MCHR1 antagonist SNAP, indicating that the effect of LC-noradrenergic neurons on rearing requires MN-derived signals. Together, these data show that, in behaving mice, LC-noradrenergic neurons exert inhibitory control over MNs and MCH-R1 dependent rearing behavior.

## DISCUSSION

We found that MNs activity acutely and reversibly increases during unsupported rearing - a frequently observed but relatively understudied behavior, whose neural triggers and modulators remain unclear despite its increasingly recognised relevance in both fundamental and translational neuroscience^26–28, 30, 33, 56^. Complementary pharmacological and optogenetic tools revealed MNs as a causal and specific driver of rearing behavior, controlling both its initiation and maintenance. Furthermore, we report *in vivo* evidence for an upstream LC→MN inhibitory signalling, which modulates rearing behavior. Together, these findings define a subcortical neural module which both tracks and controls exploratory rearing.

Our results furthermore describe several fundamental features of the relationship between MNs and rearing. Prior studies suggest that other hypothalamic neurons can exert differential control on distinct microstructural elements of self-paced behaviors (event frequency vs event duration)^57^, such as running and eating^40, 58^. This did not seem to be the case for MNs and rearing, where we observed effects on both event frequency and duration (Fig. 4A-C), suggesting that MNs orchestrate both initiation and maintenance of rearing in an MCHR-dependent manner (Fig. 4D). In our study, these effects of MNs on rearing are unlikely to be a secondary by-product of MNs’ effects on locomotion or anxiety states, since the MN manipulations that affected rearing did not affect locomotion or anxiety metrics (Fig. 2B-C, Fig. 3C). This is important, since previous studies have suggested that MNs may have an anti-locomotive effect^59–62^. However, the interpretation of chronic interventions used in these studies is complicated by compensatory effects, and more recent studies suggest that MN effects on locomotion may depend on downstream targets^63^. In our acute experiments modulating MNs, mice were fully habituated to the experimental setup before testing - to avoid suppression of rearing behavior by novelty-induced stress - and locomotion parameters remained unaltered as expected (locomotor activity Fig.3C and Fig.4D, and immobility, locomotion and turning in Fig.3G-I and Fig. 3G-I). The absence of differences in our study of changes in center/border preference in the open field - a measure of anxiety-like behavior - upon experimental manipulations of the MCH system may seem in contrast with past studies reporting that blockade of the MCHR1 receptor exerts an anxiolytic effect^64–66^. However, other studies on the role of the MCH system in anxiety-related behaviors have yielded contrasting results^67, 68^. Differences between our results and past studies on the involvement of MNs in anxiety-like behavior may be due to differences in experimental paradigms used, and further investigation will be needed to untangle the roles of MNs in exploration and anxiety. Finally, given that there is published evidence that MNs can release several neurotransmitters in addition to MCH neuropeptide^19, 43, 44^, it was important to determine whether the MN effects on rearing were mediated by MCH vs other transmitters potentially emitted by MNs. We found that rearing modulation evoked by optogenetic MN stimulation was abolished by SNAP (Fig. 4D), indicating that the SNAP-sensitive MCHR1 (the only MCH receptor in the mouse) – and thus the MCH neuropeptide – was responsible.

From the LC perspective, our results suggest both MCHR-dependent and -independent streams of LC functional output. One line of evidence suggesting this is the comparison of behavioral effects of LC manipulation in the presence and absence of SNAP (Fig. S2). Another is that, despite the LCèMN inhibitory link, we noted a dissociation between the effects of LC and MN interventions. While – among investigated behaviors – MN modulation specifically affected rearing (Figs. 2-4), modulation of LC noradrenergic neurons also affected other behaviors (Fig. S1). Interestingly, this was paralleled by a dissociation of effects of LC-noradrenergic neurons and MNs on anxiety-like behavior (Fig. S2). Following experimental manipulations of the MCH system, during self-paced behaviors in a non stressful condition no differences were observed in anxiety-like behavior, such as center avoidance in the open field (Fig. 2B, Fig. 3C). However, when the LC was activated, increased anxiety-like behavior was observed, characterized by reduced time spent in the center and reduced total distance moved (Fig. S2A-B). Conversely, inhibition of the LC significantly increased the time spent in the center (Fig. S2C). Overall, this suggests that LC-noradrenergic neurons exert wider behavioral effects than MNs, likely through projections to additional brain areas.

Our results identify important directions for future work. Since the LC is known to be activated by stress^45–48^, our findings may explain why stress reduces rearing^26^ and provides a previously unknown insight into the interplay of arousal-related LC neurons^45, 69^ and the learning and exploration-implicated MNs^12, 14, 17, 18, 22^. However, to define the role of the LCèMN circuitry in stress-induced modulation of information-gathering, it will need to be investigated in a wider range of contexts and stressors. In particular, the roles of other stress-related areas which are sources of inhibitory inputs to MNs, such as the amygdala and the BST^22^, remain to be determined. Extensive additional experiments would also be required to understand the involvement of MNs in the multiple other proposed functions of the LC, such as network resetting, brain gain control, and the inverted U relationship between arousal and performance ^47, 55, 70, 71^. Furthermore, cellular-resolution studies will be needed to assess whether rearing behavior is under control of all MNs or a specific subpopulation, to what extent that would overlap with MN activation during other awake behaviors or sleep^12–14, 72^, and what downstream circuits are involved. Our data suggests that MNs are a useful genetically-defined entry point for addressing these fundamental questions.

In summary, our study complements the increasing body of knowledge uncovering the complex and integrated roles of MNs and the lateral hypothalamus, from circuit analysis^22, 73–81^, transcriptional profiling of LH^82–84^, electrophysiology^85–89^, to behavior^12, 14, 22, 23, 72, 90^. It also adds to studies investigating naturalistic behaviors, which are proposed to improve the translational value of rodent behavioral research^56, 91^.

## METHODS

### Animal experimentation

All animal procedures were performed in accordance with the Animal Welfare Ordinance (TSchV 455.1) of the Swiss Federal Food Safety and Veterinary Office and were approved by the Zurich Cantonal Veterinary Office. Mice were kept on a standard chow and water ad libitum and on a reversed 12-h/12-h light/dark cycle. Experiments were performed during the dark phase. Adult males and females (at least 8 weeks old) were used.

### Viral vectors

The specific targeting of the GcaMP6s calcium sensor and opsins to MNs and LC-noradrenergic neurons was performed using genetic tools described and histologically validated in previous studies^12, 69^.

To target GcaMP6s to MNs, we injected an AAV vector carrying the 0.9-kb preproMCH gene promoter AAV9.pMCH.GcaMP6s.hGH (1.78 × 1014 gc/mL; Vigene Bioscience, characterized to target MCH cells with >90% specificity in ref^12^) into the lateral hypothalamus of C57BL6 mice.

To target the excitatory opsin ChrimsonR to MNs, we injected AAV-EF1a-DIO-ChrimsonR-mRuby2-KV2.1-WPRE-SV40 (5×10^11^ gc/mL; Addgene) bilaterally into the lateral hypothalamus of the previously characterized and validated MCH-Cre mice^42^, which were bred in het-WT pairs with C57BL/6 mice. Confirmation of ChrimsonR expression was performed by histology for the colocalization of mRuby and MCH staining as described previously^12^ (Fig.4A).

To target excitatory and inhibitory opsins to locus coeruleus noradrenergic neurons, we injected the Cre-dependent constructs AAV-EF1a-DIO-ChrimsonR-mRuby2-KV2.1-WPRE-SV40 (5×10^11^ gc/mL; Addgene) and AAV-EF1a-DIO-eNpHR3.0-mCherry-WPRE (5×10^12^ gc/mL; UNC Vector Core) unilaterally in the locus coeruleus of DBH-iCre mice, expressing Cre recombinase in locus coeruleus noradrenergic neurons^49^, as characterized in previous studies^92^.

For each opsin-expressing cohort of mice, corresponding control mice were littermates who underwent the same surgery, without opsin AAV injection. For every experiment, mice in each treatment group and corresponding control group were subjected to the same behavioral experimentation on the same day with a counterbalanced design.

### Stereotaxic surgery

For stereotaxic brain injections, mice were anesthetized with isoflurane and injected with Metacam (5 mg/kg of body weight, s.c.) for analgesia. In a stereotaxic frame (Kopf Instruments), a craniotomy was performed, and a 33-gauge needle mounted on a Hamilton syringe was used to inject AAV vectors.

To target the lateral hypothalamus, an injection (150 nL at a rate of 50 nL/min) was administered in one or both hemispheres (Bregma, AP −1.35 mm; ML ±0.90 mm; DV 5.30 mm; 0° angle – or Bregma, AP −1.35 mm; ML ±1.90 mm; DV 5.30 mm; 10° angle) and fibre optic implants were placed above the injection site (Bregma, AP −1.35 mm; ML, ±0.90 mm; DV, 5.00 mm; 0° angle – or Bregma, AP −1.35 mm; ML, ±1.90 mm; DV, 5.10 mm; 10° angle) based on^12, 22, 93^. To target the locus coeruleus, 2 injections (300 nL at a rate of 50 nL/min) were administered in one hemisphere (Bregma, AP -5.3 mm for females, -5.4 mm for males; ML ±0.90 mm; DV 3.7 mm, 3.4 mm) and a fibre optic implant was placed above the injection site (Bregma, AP -5.3 mm for females, -5.4 mm for males; ML ±0.90 mm; DV 3.3 mm) based on^36, 55^. Whenever unilateral targeting was used, hemispheres were counterbalanced among animals.

Before the experiments, mice were allowed to recover from surgery for at least 10 days.

### Fibre Photometry

Fibre photometry was performed using the Doric fibre photometry system, in lock-in mode using simultaneous illumination with two LEDs (405-nm and 465-nm excitation, oscillating at 334 and 471 Hz, respectively; average power, ∼100 μW at the fibre tip). Fluorescence produced by 405-nm excitation provided a real-time control for motion artifacts^94^.

### Optogenetics

The excitatory opsin ChrimsonR was activated by a red laser (635 nm; Laserglow Technologies; 30 Hz, 10 ms ON, based on^19^) and the inhibitory opsin eNpHR was activated by a yellow laser (589 nm; Laserglow Technologies; continuous wave), both yielding ∼7 mW light power at the fibre tip. The illumination protocol for ChrimsonR was 3 minutes laser OFF, 3 min laser ON, 3 min laser OFF, based on^53^. Since MN activity recovers after about 60-100 s (Fig. 2-4C), behavioral data was analysed in the first 60 seconds of ChrimsonR stimulation (Fig.6 D-F) and the baseline behavior before laser illumination was analysed in the 60 seconds before that (Fig. 5E). The illumination protocol for eNpHR was 1 min ON followed by 1.5 min OFF x 6 times based on^95, 96^, with a ramping down offset over 100 ms to avoid rebound excitation, based on^97^.

### Pharmacology

The MCH receptor antagonist SNAP-94847 hydrochloride (Tocris Bioscience, 3347) was administered IP at a dose of 20 mg/kg, dissolved in distilled water with 10% DMSO (99,5%, PanReac AppliChem, 131954.1611) and 30 mg/mL of beta-cyclodextrin (Sigma, H107), based on ref^98^. Distilled water with 10% DMSO and 30 mg/mL of beta-cyclodextrin was used as vehicle. IP administration of SNAP or vehicle was done 45 minutes prior to behavioral testing, as in ref^12^. Mice were habituated to IP injections prior to experimentation.

### Open field

Open field experiments were carried out in a 35 x 35 x 35 cm grey plexiglas box, under a ∼40 Lux illumination, to ensure a non-threatening environment favourable to the display of rearing behavior in mice^28^. Video was recorded using a camera (Basler acA1300-200um, Chromos Industrial). In Fig.5I-N, data was analysed in the time interval 3 to 6 minutes after last laser OFF. In all cases, 2 sessions were carried out for each mouse and for each condition and then averaged (except for Fig.2, where each mouse received each treatment once, and one vehicle mouse was excluded because of a technical error in IP injection, and for Fig. 5I-N, where 1 session was carried out). Mice were habituated to the apparatus before testing.

### Licking

Licking behavior was recorded in a separate chamber to avoid potential interference between self paced behaviors in the neutral environment of the open field and food-driven motivation. The test was carried out in a 19 x 19 x 35 cm plexiglas chamber installed in a ventilated, sound-insulated chest (Coulbourn Instruments) and equipped with an infrared camera (Basler acA1300-200um, Chromos Industrial), a metal spout connected to a peristaltic pump (WPM1, PeriPump) driven by an Arduino board (ArduinoUNO), and a capacitive touch sensor for the detection of licking (AT42QT1011, SparkFun Electronics). 15 µL of liquid food (strawberry milkshake) were delivered in 1-minute intervals for a total of 20 times. The output of the touch sensor was used to identify licking bouts. Each bout was defined as a cluster of consecutive licks following delivery of liquid food; bouts starting before the liquid food was made available were excluded from analysis. Mice were habituated separately to the chamber and to the milkshake in their home cage before testing.

### Fear Conditioning

The test was carried out in an operant chamber (model E10-10; Coulbourn Instruments) installed in a ventilated, sound-insulated chest and equipped with a grid floor made of stainless-steel rods (4-mm diameter). Scrambled electric shocks with a 0.5 mA intensity were delivered through the grid floor (model E13-14; Coulbourn Instruments). A tone (2.9 kHz, 90 dB, 30 sec) was delivered through intra chamber speakers. The chamber had a total floor area of 30 cm × 25 cm and a height of 29 cm, but the mouse was confined to a rectangular 17.5 cm × 13 cm region in the center, defined by a clear Plexiglas enclosure. The tone was immediately followed by a 2-seconds footshock and an ITI of 90 seconds for a total of 7 pairings.

### Video tracking and classification of behaviors in the open field

Specific behaviors were identified using a classifier based on a convolutional neural network (CNN) as in ref^40^. Briefly, the CNN was trained using >4000 movement images, generated as an RGB combining the current frame, 10 frames prior and 10 frames after (video frame rate was 30 fps), and labelled by the experimenter. The labelled frames were split between a training dataset and a validation dataset, and the training was considered complete when reaching an accuracy of ∼90% on the validation dataset, which was not used in training. The trained network was then used to classify whole experiment videos (with a sampling rate of 3 Hz). Because this tool generates motion-based RGB images, we could identify both static and dynamic behaviors. Behaviors were defined as follows: locomotion = whole body moving forward, all paws on the ground; turning = whole body rotating around the center point, all paws on the ground; immobility = body not moving, all paws on the ground, head level to the floor; grooming = body appears round and curled up, head moving, ears facing downwards; rearing = front paws are lifted, body appears shortened, head is facing up and ears backwards, no vertical surface supports the body. Supported rearing was not quantified, throughout the manuscript unsupported rearing is referred to as “rearing” for simplicity. Behavioral events were defined as uninterrupted instances of a displayed behavior. Behavioral events with a duration < 1s were excluded from analysis.

### Pupillometry

To confirm expression of the excitatory opsin ChrimsonR in LC-noradrenergic neurons, we performed a pupillometry test (Fig.5B) as in our previous work^92^. Briefly, animals were anesthetized using 2% isoflurane, their pupil was recorded with an infrared camera (20 fps) and pupil size was analysed using DeepLabCut. Optogenetic stimulation was delivered in trains of 20 Hz and 30 sec every 2-2.5 minutes.

### Data Analysis

Statistical tests and descriptive statistics were performed as specified in Results and the figure legends.

In fibre photometry experiments, to produce the plotted % ΔF/F values, the raw 405-nm–excited signal was fitted to the 465-nm–excited signal, then the % ΔF/F time series was calculated for each session as [100*(465 signal – fitted 405 signal)/fitted 405 signal], based on ref^99^. Data was z-scored to its baseline, based on^58^ (the baseline interval being -20 to -10 sec for fibre photometry data aligned with self-paced behaviors in Fig.1, and -50 to 0 sec for fibre photometry data aligned with laser onset in Fig.5; t = 0 s indicates the start of the behavior).

In Fig.1J-K, the amplitude was calculated as mean activity in the interval 0-1 seconds. In Fig.1K, using Bonferroni correction for multiple testing, p < 0.025 was considered significant. In Fig.1L each data point was obtained by calculating the maximum or minimum value in the time interval 0-1 sec for each behavioral event (using positive amplitude for behaviors associated with a positive or no deflection, and negative amplitude for those associated with a negative deflection, based on the results in Fig.1J), and averaging amplitudes for each duration bin of the behavior for each mouse. The values of p and R^2^ were calculated using Pearson’s correlation.

For behavioral data analysis, for each mouse, percent time was calculated using the formula *(100 * sum (Event Duration)) / Total Time*; event frequency was calculated as *count (Event Start) / Total Time*; event duration was calculated as *average (Event Duration)*.

Data are presented as mean ± SEM, and a P value < 0.05 was considered to indicate significance. Statistical significance was assessed using one-sample or two-sample t-test, as specified in figure legends. All t-tests were two-tailed, except for Fig. 5 D-F and I-N, where previous results (Fig. 5 C and Fig. 5 H, respectively, together with Fig. 3 and 5) led to the formulation of directional hypotheses and the use of one-tailed t-tests. All data processing and analysis was performed using custom scripts written in MATLAB R2022b (MathWorks).

## AUTHOR CONTRIBUTIONS AND ACKNOWLEDGEMENTS

This work is funded by ETH Zürich. C.C., D.P.-R., and D.B. conceived the study and designed research; S.N.D., M.P. and J.B. assisted in design of research; C.C. performed research with assistance from N.G.; C.C. and P.V. analyzed data; C.C., D.P.-R. and D.B. interpreted results; C.C., D.P.-R. and D.B. drafted the manuscript. All authors reviewed the results and contributed to and approved the final version of the manuscript. D.P.-R. and D.B. contributed equally to this work.

---

**Supplementary Figure 1:**
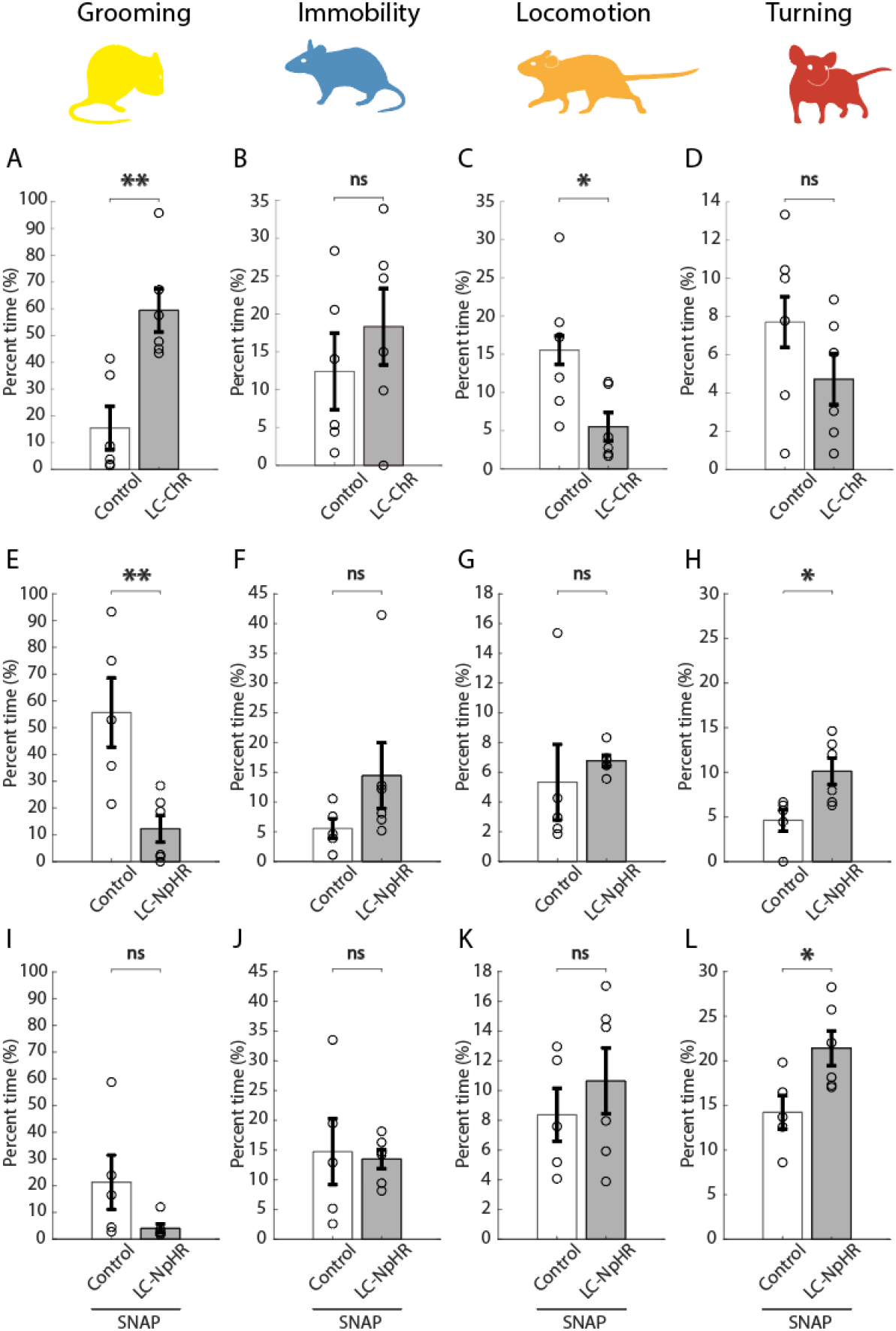
Effect of optostimulation and optoinhibition of LC neurons on other behaviors. A-D) Effect of optostimulation of LC-noradrenergic neurons on behaviors. Grooming *\*\** p = 0.0025; immobility *ns* p = 0.3933; locomotion *\** p = 0.0332; turning *ns* p = 0.2229; unpaired t-test; n = 6 LC-ChrimsonR expressing mice and 6 control mice. E-H) Effect of optoinhibition of LC noradrenergic neurons on behaviors. Grooming *\*\** p = 0.0084; immobility *ns* p = 0.1905; locomotion *ns* p = 0.5518; turning *\** p = 0.0205; unpaired t-test; n = 6 eNpHR-expressing mice and 5 control mice. I-L) Effect of optoinhibition of LC-noradrenergic neurons after injection of MCH-R1 antagonist SNAP on behaviors. Grooming *ns* p = 0.0980; immobility *ns* p = 0.8195; locomotion *ns* p = 0.4563; turning *\** p = 0.0278; unpaired t-test; n = 6 eNpHR-expressing mice and 5 control mice).

**Supplementary Figure 2:**
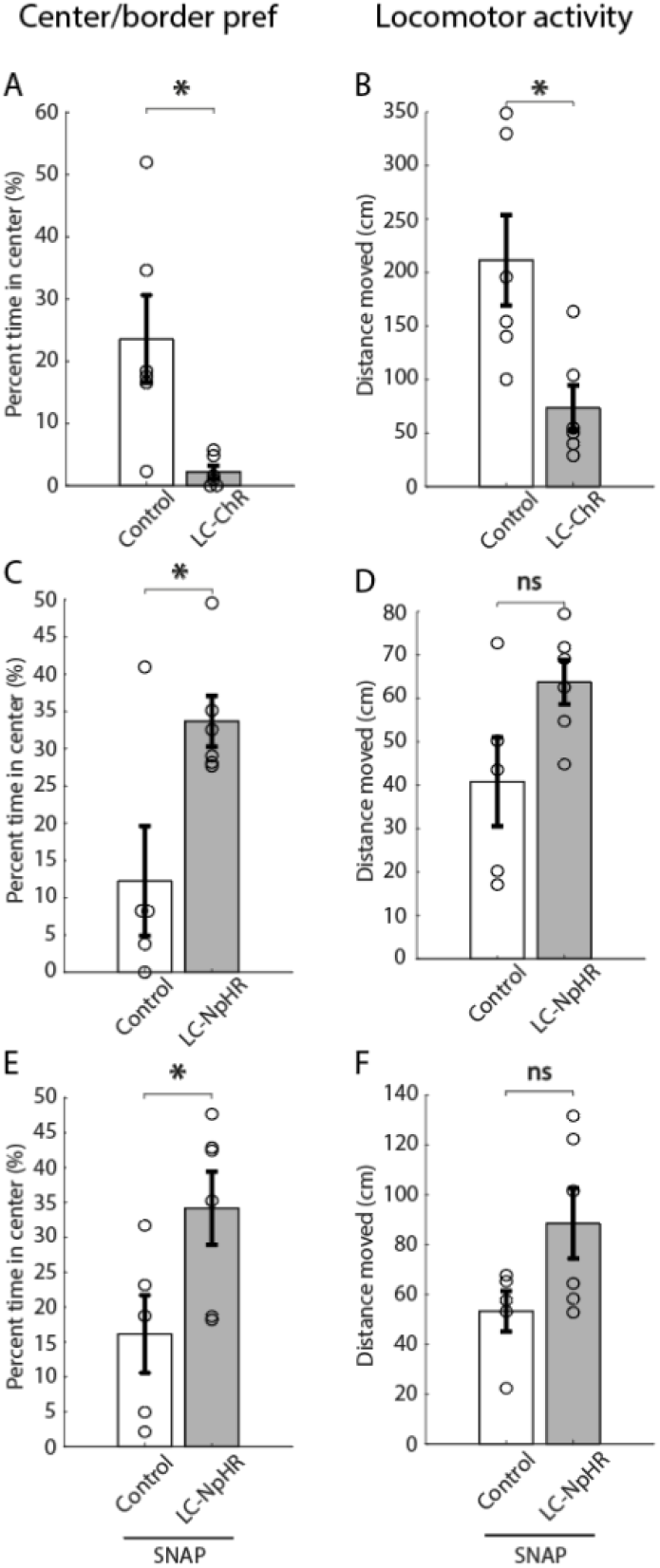
Effect of optostimulation and optoinhibition of LC neurons on center/border preference and locomotor activity in the open field. A-B) Effect of optostimulation of LC-noradrenergic neurons in control and LC-ChrimsonR mice (center preference ***\**** p = 0.0134; locomotor activity ***\**** p = 0.0153; unpaired t-test; n = 6 LC-ChrimsonR expressing mice and 6 control mice). C-D) Effect of optoinhibition of LC-noradrenergic neurons in control and LC-NpHR mice (center preference ***\**** p = 0.0202; locomotor activity ***ns*** p = 0.0629; n = 6 eNpHR-expressing mice and 5 control mice). E-F) Effect of optoinhibition of LC-noradrenergic neurons after injection of MCH-R1 antagonist SNAP in control and LC-NpHR mice (center preference ***\**** p = 0.0432; locomotor activity ***ns*** p = 0.0708; n = 6 eNpHR-expressing mice and 5 control mice).

## Notes

### Competing Interest Statement

The authors have declared no competing interest.

